# SARS-CoV-2 Spike protein binds to bacterial lipopolysaccharide and boosts proinflammatory activity

**DOI:** 10.1101/2020.06.29.175844

**Authors:** Ganna Petruk, Manoj Puthia, Jitka Petrlova, Ann-Charlotte Strömdahl, Sven Kjellström, Artur Schmidtchen

## Abstract

There is a well-known and established link between high lipopolysaccharide (LPS) levels in blood and the metabolic syndrome (MS). MS is a risk factor for developing severe COVID-19 and acute respiratory distress syndrome (ARDS). Here we define an interaction between SARS-CoV-2 Spike (S) protein and LPS and its link to aggravated inflammation in vitro and in vivo. Electrophoresis under native conditions demonstrated that SARS-CoV-2 S protein binds to *Escherichia coli* LPS, forming high molecular weight aggregates. Microscale thermophoresis analysis further defined the interaction, having a K_D_ of ~47 nM, similar to the observed affinity between LPS and the human receptor CD14. Moreover, S protein, when combined with low levels of LPS, boosted nuclear factor-kappa B (NF-κB) and cytokine responses in monocytic THP-1 cells and human blood, respectively. In an experimental model of localized inflammation, employing NF-κB reporter mice and in vivo bioimaging, S protein in conjunction with LPS significantly increased the inflammatory response when compared with S protein and LPS alone. Apart from providing information on LPS as a ligand for S protein, our results are of relevance for studies on comorbidities involving bacterial endotoxins, such as the MS, or co-existing acute and chronic infections in COVID-19 patients.

## Introduction

Coronaviruses are a group of enveloped positive-stranded RNA viruses that consist of four structural proteins including spike (S) glycoprotein, envelope (E) protein, membrane (M) protein, and nucleocapsid (N) protein (*1*). Spike glycoprotein is the most important surface protein of coronavirus including SARS-CoV-2, which can mediate the entrance to human respiratory epithelial cells by interacting with the cell surface receptor angiotensin-converting enzyme 2 (*2*). COVID-19 disease is associated with a major inflammatory component. Increased cytokine and chemokine production in response to virus infection has been the focus of several recent investigations, and patient morbidity and mortality is mainly caused by the severe systemic inflammation and acute respiratory distress syndrome (ARDS) affecting these patients (*3, 4*) although differences in ARDS disease phenotypes are noticed (*5*).

ARDS is a general systemic inflammatory reaction common for many disease states, such as pneumonia, severe infection, sepsis, burns, or severe trauma. During ARDS, activation of TLRs, such as TLR4 *via* LPS stimulation, induces an initial systemic pro-inflammatory phase characterized by a massive release of cytokines, acute phase proteins and reactive oxygen species. Additionally, activation of proteolytic cascades, like the coagulation and complement system, takes place in combination with impaired fibrinolysis, and consumption of coagulation factors and other mediators (*6–8*). Clinical symptoms of patients with ARDS therefore in many ways correspond to the pathophysiology seen during severe COVID-19 disease. There is a well-known and established link between high LPS levels in blood and MS (*9*), and obesity (*10*). Moreover, recent evidence shows that patients with MS are at risk of developing severe COVID-19 disease and ARDS. However, whether LPS plays a role in the pathogenesis of COVID-19 *per se* is at present unknown.

The above clinical and pathogenetic clues prompted us to investigate possible connections between LPS and S protein from a structural as well as functional perspective. Using electrophoresis under native conditions as well as microscale thermophoresis we indeed found that S protein binds to *Escherichia coli* LPS. S protein also boosted inflammatory responses when combined with low levels of LPS in monocytic THP-1 cells as well as in human blood. In nuclear factor-kappa B (NF-κB) reporter mice, S protein significantly increased the inflammatory response in conjunction with ultra-low, threshold levels of LPS.

## Results

### SARS-CoV-2 S protein sequence and endotoxin content

2019-nCoV full-length His-tagged S protein (R683A, R685A), composed of the S sequence Val 16 - Pro 1213 was produced in HEK293 cells and 1 μg was analyzed on SDS-PAGE followed by staining with Coomassie brilliant blue (**Fig. S1A**). The results identified a major band of ~180-200 kDa. Although the protein has a predicted molecular weight of 134.6 kDa, the result is compatible with the expected mass due to glycosylation. Next, the band was cut off from the gel and analyzed by LC-MS/MS (**Fig. S1B**). 110 peptides covered 56% of the SARS-CoV-2 S protein sequence, confirming identity. Using a limulus amebocyte lysate (LAL) assay, the LPS content in the recombinant S protein was determined to 30 fg/μg protein.

### Studies on the interaction between SARS-CoV-2 S protein and LPS

Native gel electrophoresis is used as a tool to assess structural differences in proteins, but also alterations induced by binding to external ligands. We therefore decided to study the migration of S protein alone, or in presence of increasing doses of *Escherichia coli* LPS (**Fig. 1A**). Under the conditions used, S protein migrated at the molecular mass range of 400-500 kDa. A second higher molecular 700-800 kDa band of less intensity was however observed. Addition of increasing doses of LPS indeed yielded a shift in the migration of S protein, with a reduction of particularly the 400-500 kDa band and an increase of high molecular weight material not entering the gel. Mass spectrometry of the excised protein bands was then performed. The results verified that the bands of 400-500 and 700-800 kDa were composed of S protein. S protein was also identified in the high molecular weight fraction found in the samples incubated with LPS (**Fig. 1B**). Analogously, microscale thermophoresis (MST), a highly sensitive technique probing interactions between components in solution, demonstrated interactions of fluorescence-labeled S protein with *E. coli* LPS, with a K_D_ of 46.7 ± 19.7 nM (**Fig. 1C**). For control in these experiments, we used the well-known human LPS-receptor CD14, which exhibited a K_D_ of 45.0 ± 24.3 nM to LPS. In order to gain more information on the interaction specificity, we evaluated binding of S protein to the lipid part of LPS, lipid A (**Fig. S2A**), as well as other microbial agonists (**Fig. S2B**). S protein was found to interact with lipid A (**Fig. S2A**) and also LPS from *Pseudomonas aeruginosa*, whereas no shift in the migration was observed after addition of lipoteichoic acid (LTA), peptidoglycan (PGN), or zymosan. Taken together, using two independent methods probing molecular interactions, a binding of LPS to S protein was identified. Notably, the affinity of LPS to S protein was in the range of the one observed for LPS binding to the human receptor CD14.

**Figure 1.**
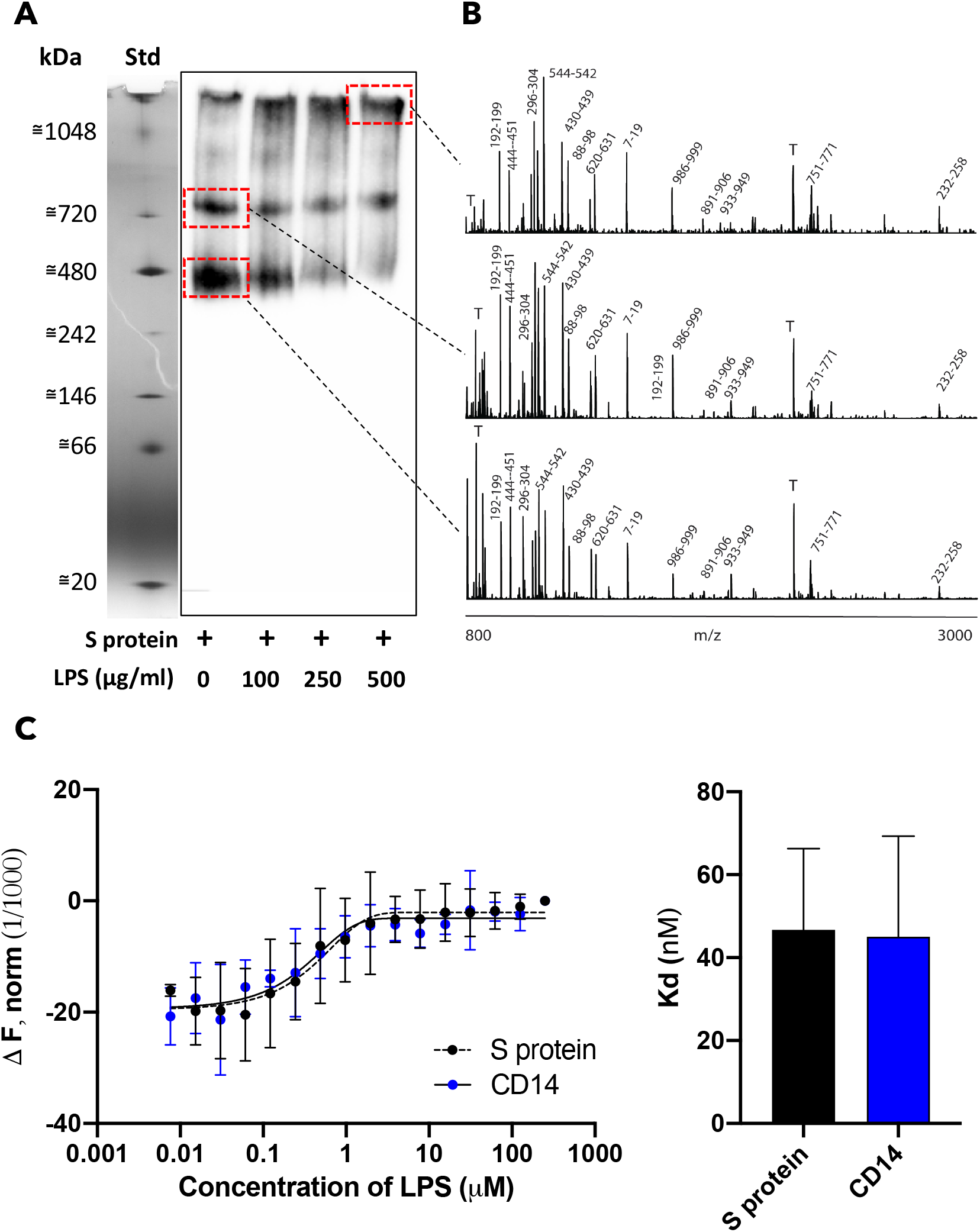
Analysis of the interaction between SARS-CoV-2 S protein and LPS *in vitro*. A) SARS-CoV-2 S protein was incubated with LPS (0-500 μg/ml), separated using Blue Native gel electrophoresis and detected by Western blot. One representative image of three independent experiments is shown (n=3). The marker lane is from the same gel but not transferred to the membrane. It is aligned and included for clarity. B) Gel pieces corresponding to the area denoted by the dotted red squares on the Western blot were cut out, in gel digestion was performed and the material was subjected to MALDI mass spectrometry analysis. Representative high resolution MALDI mass spectra are presented. The most intense tryptic fragments obtained from S protein are denoted with the sequence numbers, tryptic peptides from the autodigestion of trypsin are denoted with T. C) Microscale thermophoresis assay quantifying SARS-CoV-2 S protein interaction with LPS. CD14 was used as positive control. K_D_ constant for S protein = 46.7 ± 19.7 nM, CD14 = 45 ± 24.3 nM was determined from MST analysis. Mean ± S.D. values of six measurements are shown (n=6).

### Effects of SARS-CoV-2 S protein on LPS-induced responses in vitro

LPS effects depend on specific interactions with components of innate immunity such as LPS-binding protein (LBP), culminating in transfer of lipopolysaccharide from CD14 to Toll-like receptor 4 (TLR4) and its co-receptor MD-2 on the cell surface, leading to activation of downstream inflammatory responses (*11*). In order to probe whether the presentation and hence, activity of LPS was altered by the interaction with S protein, we decided to study the pro-inflammatory effects of S with or without LPS using THP1-XBlue-CD14 cells. After 18–24 h of incubation, NF-κB/AP-1 activation and cell metabolic activity was determined. In order to assess potential changes in the LPS-response we used a low dose of LPS of 2.5 ng/ml, which is a dose that regularly yields about 20-40% of the maximal response elicited by 100 ng/ml LPS. Addition of S protein at increasing concentrations resulted in a gradual and significant increase in NF-κB/AP-1 activation (**Fig. 2A**). It was also observed that S protein alone did not induce any significant increase in NF-κB/AP-1 activation at the concentrations used. Of relevance for the above is that the endogenous levels of LPS in the S preparation were negligible, as they were in the order of 100-1000 lower than the threshold level required for NF-κB activation. In general, patients with a systemic inflammatory response such as seen in sepsis show increased levels of LPS in plasma, with levels ranging from 0.1 to 1 ng/ml (*12*). In order to mimic those LPS levels, we therefore determined the response of the THP-1 cells to doses ranging from 0.25 ng/ml to 1 ng/ml LPS, with or without the presence of 5 nM of S protein. It was observed that NF-κB activation was significantly boosted even at those low doses of LPS. Notably, LPS at 0.25 ng/ml, which alone did not induce a significant increase of NF-κB activation, yielded a significant response together with S protein. It was also observed that LPS at doses of 0.5-1 ng/ml, combined with S protein, yielded response levels produced by 10 ng/ml LPS (**Fig. 2B**). In these studies, cell viability was regularly measured by the MTT assay, and no significant toxic effects were detected (**Fig. 2A, B**). Finally, using human blood, we observed a similar increase of the LPS response. Again, particularly ultralow levels of LPS, 50 pg/ml, showed boosted TNF-α levels together with S protein (**Fig. 2C**). Taken together these results unequivocally demonstrate that S protein increases LPS responses in vitro in monocytic cells and human blood, and in particular, that the activation seen by low, threshold levels of LPS is boosted several-fold by the addition of S protein.

**Figure 2.**
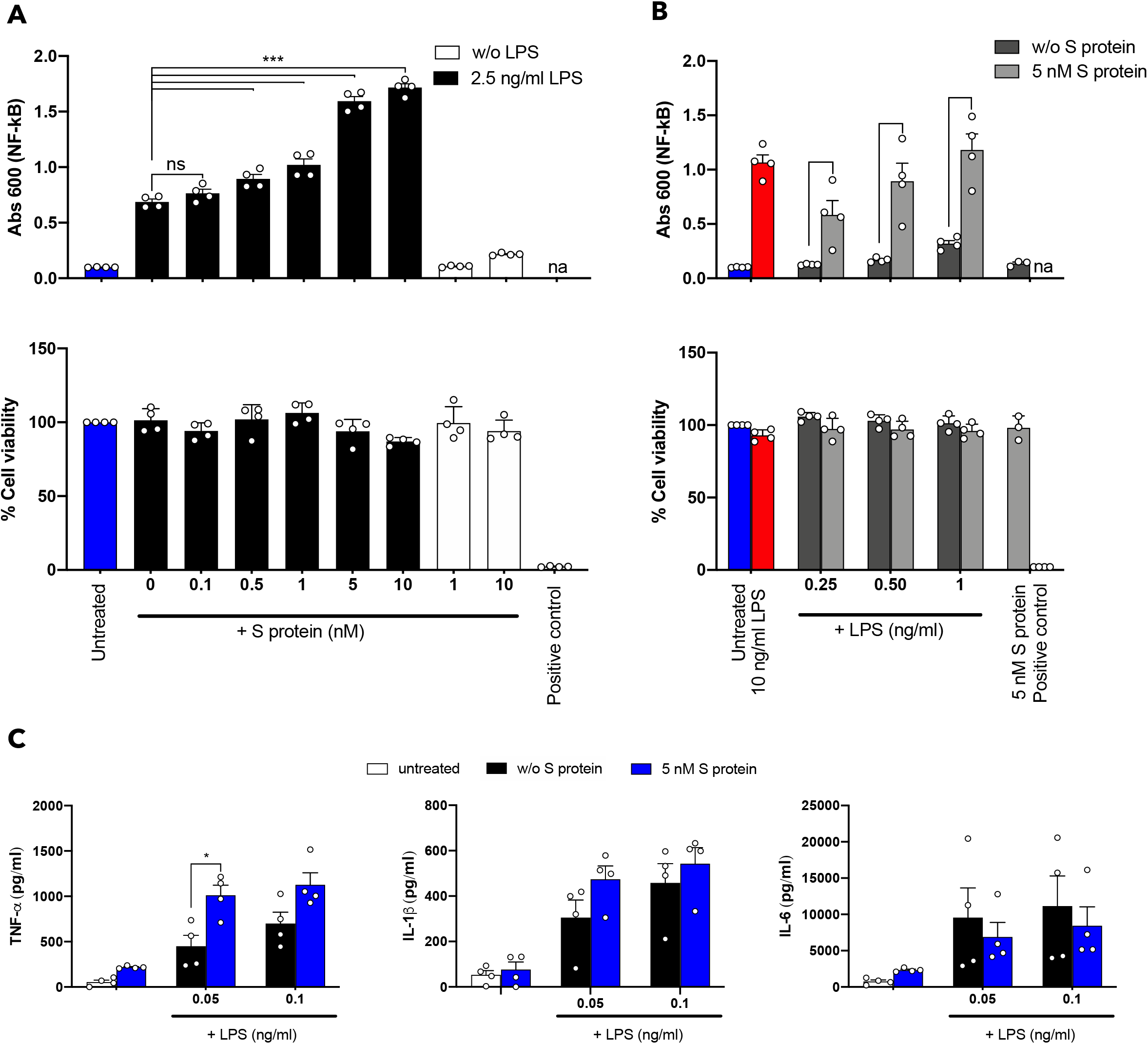
Effects of SARS-CoV-2 S protein on LPS-induced responses in THP-1 cells. THP-1-XBlue-CD14 cells were treated with increasing concentrations of SARS-CoV-2 S protein (0-10 nM) and a constant dose of LPS (2.5 ng/ml) (A) or with increasing doses of LPS (0.25-1 ng/ml) and constant amount of S protein (5 nM) (B). MTT viability assay for analysis of toxic effects of S protein and LPS on THP-1 cells is shown in lower panels for A and B. C) Cytokine analysis of blood collected from healthy donors, 24 h after treatment with S protein with or without 0.05 and 0.1 ng/ml LPS. Untreated blood was used as a control. The mean ± S.E.M. (NF-kB and blood assays) or S.D. (MTT assay) values of four independent experiments performed in duplicate are shown. *, p< 0.05; ***, p< 0.001; ****, p < 0.0001, determined using two-way ANOVA with Sidak’s multiple comparisons test (NF-kB and blood assays) or one-way ANOVA with Dunnett’s multiple comparison test (MTT assay). na; not analyzed, w/o; without.

### Effects of SARS-CoV-2 S protein on endotoxin responses in an experimental mouse model

In an experimental animal model, we wanted to simulate a situation of localized endotoxin induced inflammation. In previous models, we utilized doses of 100 μg LPS injected subcutaneously, a dose level which yielded a robust and significant LPS response (*13*). In this modified model, similar to the strategy described above on the THP-1 cells, we employed low, threshold levels comprising 2 μg LPS which were injected subcutaneously with or without 5 μg S protein. Using mice reporting NF-κB activation, we indeed found that the addition of S protein significantly increased the inflammatory response (**Fig. 3**). S alone at the dose of 5 μg did not yield any significant inflammatory response. Apart from a strongly increased response by the LPS and S combination, we also observed that the LPS-S protein mix resulted in a prolonged inflammatory response. Taken together, the results demonstrated that SARS-CoV-2 S protein also retains its boosting effect in conjunction with LPS in a subcutaneous model of endotoxin-driven inflammation.

**Figure 3.**
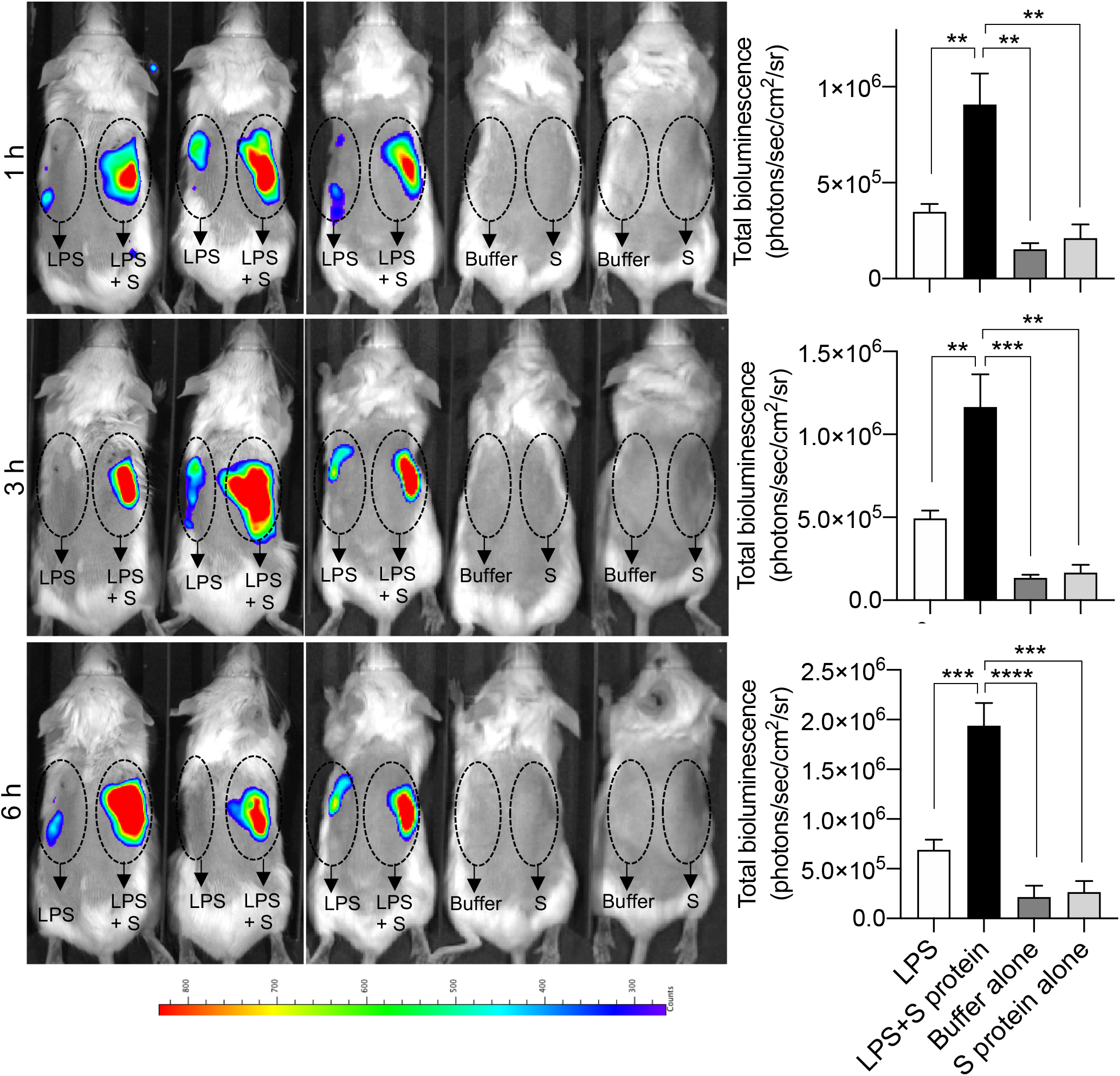
SARS-CoV-2 S protein combined with LPS boosts inflammation in NF-κB reporter mice. *In vivo* inflammation imaging in NF-κB reporter mice. LPS alone or in combination with SARS-CoV-2 S protein (S) was subcutaneously deposited on the left and right side, respectively, on the back of transgenic BALB/c Tg(NF-κB-RE-luc)-Xen reporter mice. Non-invasive *in vivo* bioimaging of NF-κB reporter gene expression was performed using the IVIS Spectrum system. Representative images show bioluminescence at 1, 3 and 6 h after subcutaneous deposition. A bar chart shows measured bioluminescence intensity emitted from these reporter mice. Dotted circles represent area of subcutaneous deposition and region of interest for data analysis. Data are presented as the mean ± SEM (*n* = 5 mice for LPS group, 5 mice for LPS and S protein group, 3 mice for buffer control, and 3 mice for S protein control). *P* values were determined using a one-way ANOVA with Holm-Sidak posttest. \*\**P* ≤ 0.01; \*\*\**P* ≤ 0.001; *\*\*\*\*P ≤* 0.0001; NS, not significant.

## Discussion

Here we demonstrate a previously unknown interaction between SARS-CoV-2 S protein and LPS, leading to a boosting of pro-inflammatory actions *in vitro* as well as *in vivo*. These results on the synergism between LPS and S protein are of clinical importance, as this could give new insights in the comorbidities that may increase the risk for severe COVID-19 disease and ARDS, and its pathogenetic steps.

As mentioned in the Introduction, the observed link between high LPS levels in blood and the MS (*9*), and the fact that MS is a risk factor for developing severe COVID-19 prompted this study (*1*). However, as summarized in **Table 1**, the clinical implications may be broader and go beyond MS. Notable, in patients with chronic obstructive pulmonary disease (COPD), COVID-19 infection is associated with substantial severity and mortality rates (*14*). Therefore, a causative link between LPS derived from bacterial colonization and infection of the lungs in COPD patients and COVID-19 severity could also be proposed here. Related to this, is the finding that there is a correlation between LPS levels and bacterial loads during pneumonia (*15*). Moreover, compared to former and never smokers with COPD, current smokers are at greater risk of severe COVID-19 complications and higher mortality rate (*14*), and intriguingly, bacterial LPS is an active component of cigarette smoke (*16*). Increased endotoxin levels are also observed in patients with inflammatory bowel disease (IBD) (*17*). The observations that all these comorbidities are risk factors for severe COVID-19 lend further support for a pathogenetic link to SARS-CoV-2 infection and endotoxinemia, Moreover, intriguingly, Kawasaki disease in children, which has been reported in young COVID-19 patients (*18*) as well as in patients with SARS HCoV-NH (*19*) has been linked to LPS as a trigger (*20*). Other possible comorbidities that should be considered is periodontitis, where LPS from *Porphyromonas gingivalis* and other bacteria can reach the systemic circulation (*21*). Indeed, a recent hypothesis on this matter has been raised (*22*). All these observations on links between LPS levels and several diseases and conditions (**Table 1**), and the risk for developing severe COVID-19 (*1*) implies that measurement of endotoxin levels in COVID-19 patients could have significant diagnostic implications and be of relevance for patient management and treatment decisions. Clearly, clinical prospective studies are mandated in order to assess whether the findings from the present study can be translated to the clinical setting.

**Table I.** Diseases involving endotoxins and links to COVID-19

| **Disease/indication** | **LPS levels in plasma or serum** | **Ref** | **Bacterial influence** | **Ref** | **COVID-19 and risk for severe disease** | **Ref** |
| --- | --- | --- | --- | --- | --- | --- |
| **Metabolic syndrome (MS)** | 47 - 96 EU/ml | (9, 31, 32) | Dysbiosis | (33) | 12% | (34, 35) |
| **Chronic obstructive pulmonary disease (COPD)** | Unknown | (36) | Microbial colonization | (37, 38) | 1-3% | (14) |
| **Inflammatory bowel syndrome (IBD)** | 44.41 ± 89.44 pg/ml | (17, 39) | Dysbiosis | (40, 41) | Unknown | (42) |
| **Kawasaki disease** | Unknown | (43) | Unknown | - | Unknown | (44) |
| **Periodontitis** | 0.89 ± 2.90 ng/ml | (45, 46) | Microbial colonization | (47) | Unknown | (22) |

Although the here disclosed binding of LPS to SARS-CoV-2 S protein is novel, the interaction between S proteins and endotoxins is not necessarily new to nature. Indeed, interactions between viruses and bacteria for the induction of severe respiratory disease have been described since the early thirties (*23*). Obviously, multiple complex and diverse inflammatory mechanisms may underlie this general finding. However, it is worth noting that recent observations from porcine animal models indeed demonstrated that infection with porcine respiratory coronavirus, a highly prevalent virus in swine populations significantly sensitizes the lungs to LPS (*24*). Notably, the effects of separate virus or LPS inoculation were subclinical and failed to induce sustained cytokine levels, whereas the combination of the two agents significantly triggered severe respiratory disease and enhanced particularly TNF-α levels (*25*), findings indeed relevant in the light of the present results showing boosted TNF-α levels in human blood with the combination of LPS and SARS-CoV-2 S protein. In agreement with this, it is also worth noting that the disease denoted “Shipping fever”, which affects cattle particularly in relation to stress and transports, can be triggered by a combination of bovine respiratory corona virus (BCoV) and inhaled LPS (*26*).

The exact molecular mechanism underlying the observed boosting of inflammation by SARS-CoV-2 S protein remains to be investigated. Clearly, MST analysis measuring the interaction between S protein and LPS yielded a K_D_ in the nM range, which indicates a high-affinity binding. Moreover, electrophoresis under native conditions confirmed the interaction, and also showed that higher molecular weight S protein aggregates were formed upon this interaction. Notably, such aggregates were also induced by lipid A, the core of the LPS molecule, common for endotoxin producing bacteria. Previous reports indicate that both LPS and protein aggregates can be proinflammatory (*27, 28*), and moreover, the well-known LPS-binding protein (LBP) forms proinflammatory aggregates with LPS particularly at a high LPS/LBP ratio (*29*). It is therefore possible that LPS-triggered oligomerization or aggregate formation by SARS-CoV-2 S protein help explain the observed boosting of inflammation. In addition, the observation that LPS showed a similar affinity to S protein as for CD14, implies that S protein interference with the normal presentation and clearance of LPS induces a dysfunctional inflammatory state.

In conclusion, we report a previously undisclosed interaction between the Spike protein of SARS CoV-2 and LPS, and its link to induction of NF-κB and cytokine responses in monocytes and human blood, as well as increased NF-κB responses in experimental animal models. The interaction between S protein and LPS therefore provides a new therapeutic target enabling development of drugs that can ameliorate the hyperinflammation seen during COVID-19 infection.

## Materials and Methods

### Proteins

SARS-Cov-2 S protein was produced by ACROBiosystems (USA). The sequence contains AA Val 16 - Pro 1213 (Accession # QHD43416.1 (R683A, R685A)). Briefly, the 2019-nCoV Full Length S protein (R683A, R685A), His Tag (SPN-C52H4) was expressed in human 293 cells (HEK293) and purified. The protein was lyophilized from a 0.22 μm filtered solution in 50 mM Tris, 150 mM NaCl, pH7.5. Lyophilized product was reconstituted in endotoxin free water, aliquoted and stored at −80 °C according to the manufacturer’s protocol. The purity was >85%. Human His-Tag-CD14 (hCD14-his) was produced recombinantly in insect cells by using the Baculovirus Expression Vector System (BEVS). Since this construct is secreted, media was centrifuged in a JLA8-1000 rotor at 8000 g, 20 min, 4°C and then the supernatant was filtered with a PES 0.45 μm filter top (0.45 μm pore size). Subsequently hCD14-his was purified on a 5 mL HisTrap Excel column (GE Healthcare) by employing ÄKTA Pure system (GE Healthcare). Eluted fractions were analyzed by precast SDS-PAGE gel (Bio-Rad) stained with BioSafe Coomassie (Bio-Rad) or subjected to Western blot. Peak fractions containing the protein of interest were pooled and digested with tobacco etch virus (TEV) protease to remove the His-Tag. After TEV digestion, the protein solution was run a second time on the His-Trap column. Fractions containing the protein were collected, pooled and purified further on a HiLoad 26/60 Superdex 75 pg gel filtration column. At the end of purification, the purity of hCD14 was estimated to >90%. The protein was aliquoted and stored at −80 °C before use.

### Limulus amebocyte lysate (LAL) assay

The content of endotoxin in 1 μg purified SARS-CoV-2 S protein was analyzed using a commercially available Pierce™ Chromogenic Endotoxin Quant Kit (Thermo-Fisher, USA), according to the manufacturer’s protocol with small modifications. In particular, the standard curve was done with lipopolysaccharide (LPS) from *E. coli* (Sigma, USA) in the range between 0.01-10 pg/ml. All samples were prepared in endotoxin-free tubes kept in a thermoblock set to 37 °C. At the end of the incubation, 150 μl of each sample were transferred to 96-wells plates and analyzed for absorbance at 405 nm using a spectrophotometer. Pyrogen-free water, used to dissolve the protein, was used as negative control.

### SDS-PAGE

1 μg of SARS-Cov-2 S protein was diluted in loading buffer and loaded on 10–20% Novex Tricine pre-cast gel (Invitrogen, USA). The run was performed at 120 V for 1 h. The gel was stained by using Coomassie Brilliant blue (Invitrogen, USA). The image was obtained using a Gel Doc Imager (Bio-Rad Laboratories, USA).

### Blue Native (BN)-PAGE

2 μg of SARS-Cov-2 S protein were incubated with 0.1, 0.25 or 0.5 mg/ml of *E. coli* LPS or Lipid A for 30 min at 37 °C in 20 μL as final volume. At the end of the incubation the samples were separated under native conditions on BN-PAGE (Native PAGE BisTris Gels System 4–16%, Invitrogen) according to the manufacturer’s instructions. Proteins were visualized by Coomassie staining. For Western blotting, the material was subsequently transferred to a PVDF membrane using the Trans-Blot Turbo (Bio-Rad, USA). Primary antibodies against the His-tag (1:2000, Invitrogen) were followed by secondary HRP conjugated antibody (1:2000, Dako, Denmark), for detection of S protein. The protein was visualized by incubating the membrane with SuperSignal West Pico Chemiluminescent Substrate (Thermo Scientific, Denmark) for 5 min followed by detection using a ChemiDoc XRS Imager (Bio-Rad). In another set of experiments, 2 μg of SARS-Cov-2 S protein were incubated with 0.25 mg/ml of LPS and Lipid A from *E. coli*, LPS from *P. aeruginosa*, LTA and PGN from *S. aureus*, and zymosan from *S. cerevisiae*. BN-PAGE and Western blotting were performed as described above. LPS from *E. coli* and *P. aeruginosa* as well as Lipid A were purchased from Sigma-Aldrich, whereas LTA, PGN and zymosan were purchased from InvivoGen.

### Mass Spectrometry analysis

After separation by SDS- or BN-PAGE and Coomassie staining, bands in the gels were cut out and the digestion was performed according to Shevchenko et al. (*30*). Briefly the gel pieces were washed with water, then with a mix of 50 mM ammonium carbonate in 50% acetonitrile (ACN). Gel pieces were shrunk with 100% of acetonitrile and then reduced with 10 mM DTT 30 min 56 °C. Alkylation was performed with 55 mM idoacetamide at RT. 10 ng/μl of trypsin solution was added to cover the gel-pieces placed on ice and after 1 hour the samples were placed at 37 °C and overnight digestion was performed. The supernatant was acidified using 5% of formic acid and then analyzed by MALDI MS or LC-MS/MS.

### MALDI mass spectrometry analysis

For MALDI mass spectrometry analysis, digested SARS-CoV-2 S protein samples were mixed with a solution of 0.5 mg/ml α-Cyano-4-hydroxycinnamic acid (CHCA) 50% ACN/0.1 % phosphoric acid solution directly on a stainless MALDI target plate. Subsequent MS analysis was performed on a MALDI LTQ Orbitrap XL mass spectrometer (ThermoScientific, Bremen, Germany). Full mass spectra were obtained by using the FT analyser (Orbitrap) at 60,000 resolution (at *m/z* 400). Recording of Mass spectra was performed in positive mode with a 800–4000 Da mass range. The nitrogen laser was operated at 27 μJ with automatic gain control (AGC) off mode using 10 laser shots per position. Evaluation of the spectra was performed with Xcalibur v 2.0.7. software (from Thermo Fisher Scientific, San José, CA).

### LC-MS/MS

The LC-MS/MS detection was performed on HFX orbitrap equipped with a Nanospray Flex ion source and coupled with an Ultimate 3000 pump (Thermo Fischer Scientific). Peptides were concentrated on an Acclaim PepMap 100 C18 precolumn (75 μm x 2 cm, Thermo Scientific, Waltham, MA) and then were separated on an Acclaim PepMap RSLC column (75 μm x 25 cm, nanoViper, C18, 2 μm, 100 Å) with heating at 45 °C for both the columns. Solvent A (0.1% formic acid) and solvent B (0.1% formic acid in 80% ACN) were used to create a nonlinear gradient to elute the peptides. For the gradient, the percentage of solvent B increased from 4% to 10% in 20 min, increased to 30% in 18 min and then increased to 90% in 2 min and kept it for a further 8 min to wash the columns.

The Orbitrap HFX instrument was operated in data-dependent acquisition (DDA) mode. The peptides were introduced into mass spectrometer *via* stainless steel Nanobore emitter (OD 150 μm, ID 30 μm) with the spray voltage of 1.9 kV and the capillary temperature was set 275 °C. Full MS survey scans from m/z 350-1600 with a resolution 1200,000 were performed in the Orbitrap detector. The automatic gain control (AGC) target was set to 3 × 10^6^ with an injecting time of 20 ms. The most intense ions (up to 20) with charge state 2-5 from the full scan MS were selected for fragmentation in Orbitrap. MS2 precursors were isolated with a quadrupole mass filter set to a width of 1.2 m/z. Precursors were fragmented by collision-induced dissociation (CID) with a collision energy of 27%. The resolution was set at 15000 and the values for the AGC target and inject time were 2 × 10^3^ and 60 ms, respectively for MS/MS scans. The duration of dynamic exclusion was set 15 s and the mass tolerance window 10 ppm. MS/MS data was acquired in centroid mode. MS/MS spectra were searched with PEAKS (version 10) against UniProt Homo Sapiens (version 2020_02). A 10 ppm precursor tolerance and 0.02 Da fragment tolerance were used as the MS settings. Trypsin was selected as enzyme with one missed cleavage allowance, methionine oxidation and deamidation of aspargine and glutamine were treated as dynamic modification and carbamidomethylation of cysteine as a fixed modification. Maximum of post-translational modifications (PTM) per peptide was 2.

### Microscale thermophoresis

Microscale thermophoresis (MST) was performed on a NanoTemper Monolith NT.115 apparatus (Nano Temper Technologies, Germany). 40 μg SARS-CoV-2 S protein and 100 μg of recombinant hCD14 were labeled by Monolith NT Protein labelling kit RED – NHS (Nano Temper Technologies, Germany) according to the manufacturer’s protocol. 5 μl of 20 nM labeled SARS-CoV-2 S protein or 20 nM labeled hCD14 were incubated with 5 μl of increasing concentrations of LPS (250–0.007 μM) in 10 mM Tris pH 7.4. Then, samples were loaded into standard glass capillaries (Monolith NT Capillaries, Nano Temper Technologies) and the MST analysis was performed (settings for the light-emitting diode and infrared laser were 80%). Results shown are mean values ± SD of six measurements.

### NF-κB activation assay

THP1-XBlue-CD14 reporter cells (InvivoGen, San Diego, USA) were seeded in 96 well plates in phenol red RPMI, supplemented with 10% (v/v) heat-inactivated FBS and 1% (v/v) Antibiotic-Antimycotic solution (180,000 cells/well). Cells were treated with 2.5 ng/ml LPS (Sigma, USA) with increasing concentrations (0.1-10 nM) of SARS-CoV-2 S protein or with 5 nM SARS-CoV-2 S protein with increasing concentrations of LPS (0.25-1 ng/ml). Then, the cells were incubated at 37 °C for 20 h. At the end of incubation, the NF-κB activation was analyzed according to the manufacturer’s instructions (InvivoGen, San Diego, USA), i.e. by mixing 20 μl of supernatant with 180 μl of SEAP detection reagent (Quanti-BlueTM, InvivoGen), followed by absorbance measurement at 600 nm. Data shown are mean values ± SEM obtained from at least four independent experiments all performed in triplicate.

### MTT assay

The cytotoxicity of the treatments was evaluated by adding 0.5 mM Thiazolyl Blue Tetrazolium Bromide to the cells remaining from NF-κB activation assay. After 2 h of incubation at 37 °C, cells were centrifuged at 1000 g for 5 min and then the medium was removed. Subsequently, the formazan salts were solubilized with 100 μl of 100 % DMSO (Duchefa Biochemie, Haarlem). Absorbance was measured at a wavelength of 550 nm. Cell survival was expressed as percentage of viable cells in the presence of different treatment compared with untreated cells. Lysed cells were used as positive control. Data shown are mean values ± SD obtained from at least four independent experiments all performed in triplicate.

### Blood assay

Fresh venous blood was collected in the presence of lepirudin (50 mg/ml) from healthy donors. The blood was diluted 1:4 in RPMI-1640-GlutaMAX-I (Gibco) and 1 ml of this solution was transferred to 24-well plates and stimulated with 0.05 or 0.1 ng/ ml of LPS in the presence or the absence of 5 nM of SARS-CoV-2 S protein. After 24 h incubation at 37 °C in 5% CO2, the plate was centrifuged for 5 min at 1000 *g* and then the supernatants were collected and stored at −80 °C before analysis. The experiment was performed at least 4 times by using blood from different donors each time.

### ELISA

The cytokines tumor necrosis factor alpha (TNF-α), interleukin-1β (IL-1β) and interleukin-6 (IL-6), were measured in human plasma obtained after the blood experiment described above. The assay was performed by using human inflammation DuoSet^®^ ELISA Kit (R&D Systems) specific for each cytokine, according to the manufacturer’s instructions. Absorbance was measured at a wavelength of 450 nm. Data shown are mean values ± SEM obtained from at least four independent experiments all performed in duplicate.

### Mouse inflammation model

The immunomodulatory effects of 5 μg SARS-CoV-2 S protein in combination or not with 2 μg LPS/mouse were tested employing BALB/c tg (NF-B-RE-Luc)-Xen reporter mice (Taconic Biosciences, Albany, NY, USA, 10–12 weeks old). The dorsum of the mouse was shaved carefully and cleaned. SARS-CoV-2 S protein was mixed with LPS immediately before subcutaneous injection on the dorsums of the mice anesthetized with isoflurane (Baxter, Deerfield, IL, USA). Then the animals were transferred to individually ventilated cages and imaged at 1, 3, and 6 h after the injection. An In Vivo Imaging System (IVIS Spectrum, PerkinElmer Life Sciences) was used for the longitudinal determination of NF-κB activation. Fifteen minutes before the IVIS imaging, mice were intraperitoneally given 100 μl of D-luciferin (150 mg/kg body weight). Bioluminescence from the mice was detected and quantified using Living Image 4.0 Software (PerkinElmer Life Sciences).

### Ethics statement

All animal experiments are performed according to Swedish Animal Welfare Act SFS 1988:534 and were approved by the Animal Ethics Committee of Malmö/Lund, Sweden.

### Statistical analysis

All in vitro assays were repeated at least three times. Unless otherwise stated. Data are presented as means ± SEM. Differences in the mean between two groups were analyzed using Student’s *t* test for normally distributed data and Mann-Whitney test otherwise. To compare means between more than two groups, a one-way ANOVA with Dunnet or Holm-Sidak posttest were used. Statistical analysis, as indicated in each figure legend, were performed using GraphPad Prism software v8. P values <0.05 were considered to be statistically significant.

## Acknowledgements

This work was supported by grants from the Swedish Research Council (project 2017-02341), the Welander-Finsen, Crafoord, and Österlund Foundations, The Royal Physiographic Society of Lund, and The Swedish Government Funds for Clinical Research (ALF).

## Abbreviations

ARDS: acute respiratory distress syndrome
COVID-19: coronavirus disease 2019
MS: metabolic syndrome
LBP: LPS-binding protein
LPS: lipopolysaccharide
NF-κB: nuclear factor-kappa B
SARS-CoV-2 Spike protein: S protein
TLR4: Toll-like receptor 4

---

**Figure S1.**
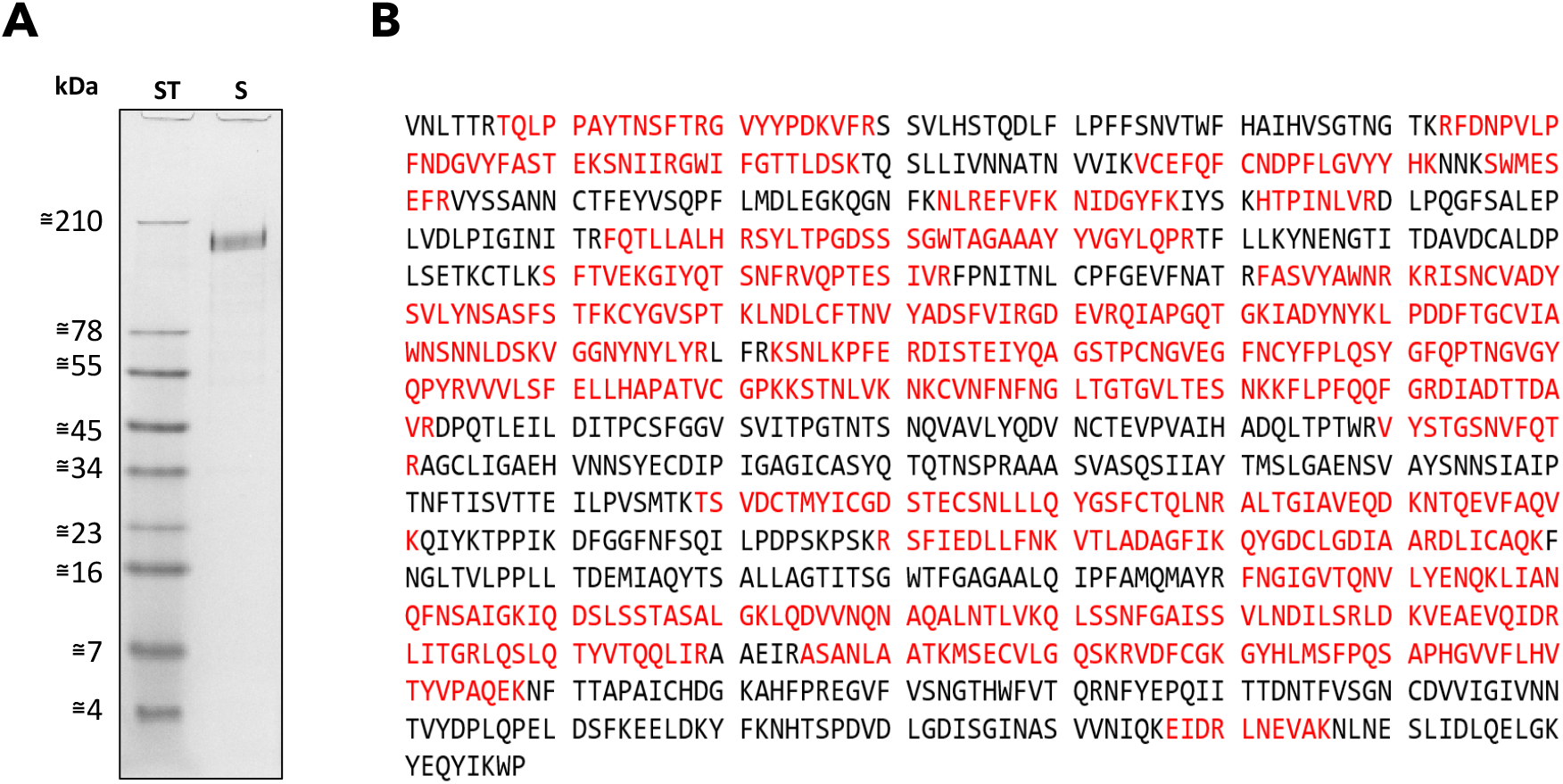
SARS-CoV-2 S protein sequence and endotoxin content. A) 1 μg of SARS-CoV-2 S protein was separated by SDS-PAGE (16.5% Tris-Tricine gel) followed by Coomassie staining. B) LC-MS/MS data were obtained after in gel digestion of SARS-CoV-2 S protein after SDS-PAGE separation. Database analysis confirmed the identity of the recombinant protein to SARS-CoV-2 S protein identifying 56% of the protein sequence as shown in red. Totally 110 peptides correspond to the protein sequence, two other proteins were detected i.e. keratin type II (25% sequence coverage, 14 unique peptides) and keratin type I (9% sequence coverage, 5 unique peptides.)

**Figure S2.**
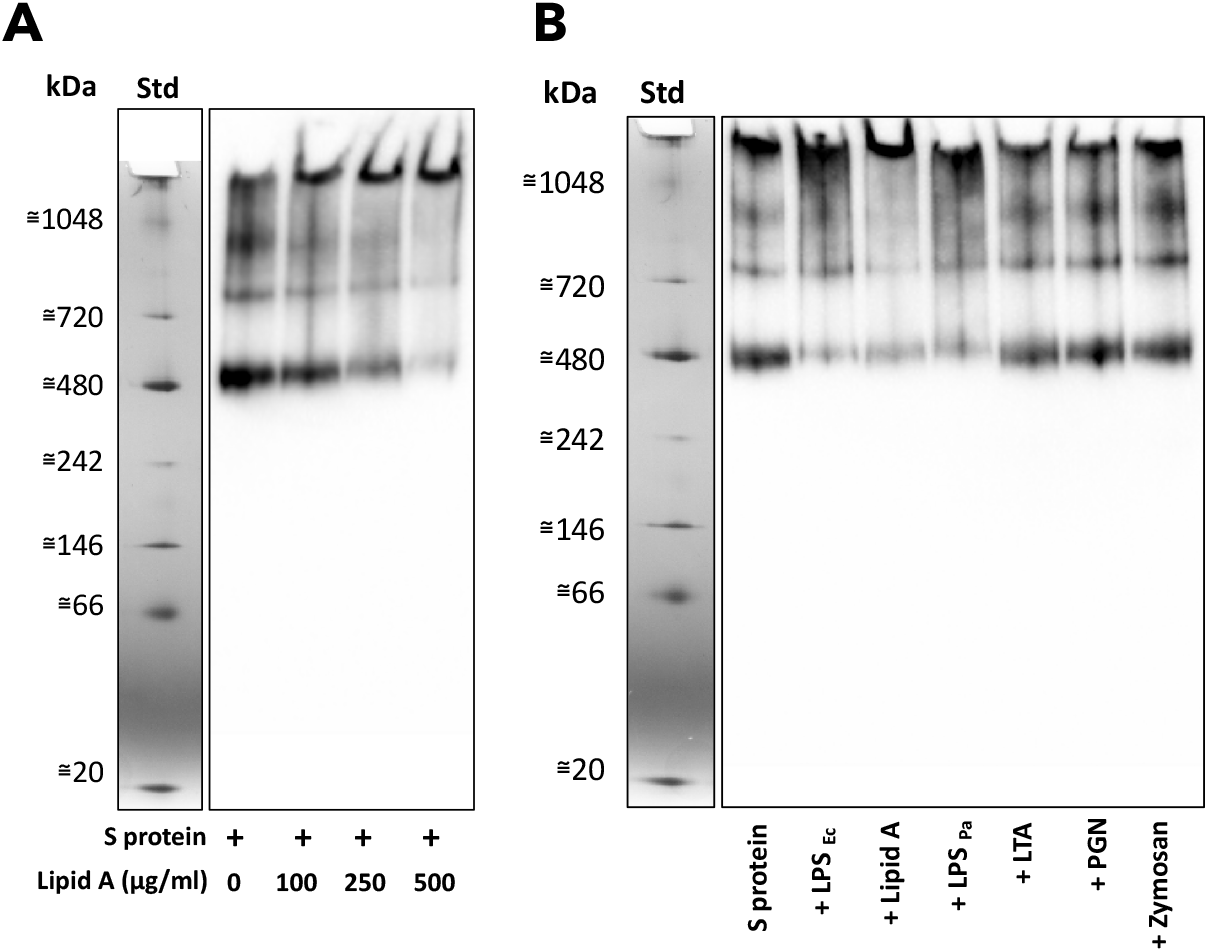
Analysis of binding of SARS-CoV-2 S protein to different TLR ligands. BN-PAGE followed by Western blotting of S protein incubated with A) 0-0.5 mg/ml of Lipid A, or B) with 0.25 mg/ml of LPS from *E. coli* (LPS _Ec_), Lipid A, LPS from *P. aeruginosa* (LPS _Pa_), LTA, PGN and zymosan.

